# Three-dimensional chromatin re-organization during muscle stem cell aging

**DOI:** 10.1101/2022.09.21.508601

**Authors:** Benjamin A. Yang, Jacqueline A. Larouche, Kaitlyn M. Sabin, Paula M. Fraczek, Stephen C. J. Parker, Carlos A. Aguilar

## Abstract

Age-related skeletal muscle atrophy or sarcopenia is a significant societal problem that is becoming amplified as the world’s population continues to increase. A critical contributor to sarcopenia is the loss in the number and function of muscle stem cells, which maintain tissue homeostasis and regenerate damage. The molecular mechanisms that govern muscle stem cell aging encompass changes across multiple regulatory layers and are integrated by the three-dimensional organization of the genome. To quantitatively understand how hierarchical chromatin architecture changes during muscle stem cell aging, we generated 3D chromatin conformation maps (Hi-C) and integrated these datasets with multi-omic (chromatin accessibility and transcriptome) profiles from bulk populations and single cells. We observed that muscle stem cells display static behavior at global scales of chromatin organization during aging and extensive rewiring of local contacts at finer scales that were associated with variations in transcription factor binding and aberrant gene expression. These data provide insights into genome topology as a regulator of molecular function in stem cell aging.

## INTRODUCTION

The growing population over the age of 65 presents significant healthcare challenges^1^ including reduced mobility and increased frailty, which is nominally associated with declines in the volume, health and repair of skeletal muscle. Skeletal muscle is regenerated by a population of resident muscle stem cells (MuSCs)^2,3^ that decrease in number and function with age^4,5^. The molecular mechanisms that govern MuSC dysfunction in old age encompass changes across multiple inter-connected molecular systems^6–8^ which collectively converge to drive aberrant chromatin packaging and three-dimensional folding of the genome^9^. 3D genomic packaging of chromatin plays a primary role in regulating cellular functions by physically contacting distal regulatory elements such as enhancers with genes^10^, facilitating changes in transcription^11^, yet how this architecture is modified in MuSCs in old age remains unknown^12^.

To probe how old age induces changes in nuclear organization in MuSCs, we performed *in situ* chromosome conformation capture followed by sequencing (Hi-C) on MuSCs isolated from young and aged skeletal murine muscles. We integrated the Hi-C maps with gene expression (RNA-seq), chromatin modifications associated with promoters (H3K4me3), and chromatin accessibility (ATAC-seq)^5^ profiles, and observed stable chromatin architecture at the level of chromatin compartments. These results contrasted with topologically associated domains (TADs) and chromatin loops that displayed dynamic restructuring and encompassed genes associated with cell cycle regulation, maintenance of quiescence, and cellular stress. To increase the resolution of our approach, we generated single-cell ATAC-seq profiles of young and aged MuSCs and integrated these datasets with age-matched single-cell RNA-seq datasets. This approach revealed extensive rewiring within chromatin hubs at the level of enhancer-promoter contacts that was linked to alterations in gene expression. Together, this work represents a rich multi-omic framework that provides insights into the regulation of pathological gene expression programs that attenuate MuSC regenerative potential in old age.

## RESULTS

### Profiling of Global 3D Genome Organization in Muscle Stem Cells During Aging

To understand how MuSC genome organization is modified in aging, hind limb muscles (tibialis anterior, gastrocnemius, extensor digitorum longus, quadriceps) were isolated from young (3 months) and aged (24-26 months) mice. Fluorescent activated cell sorting (FACS) with both negative (Sca-1^−^, CD45^−^, Mac-1^−^, Ter-119^−,^ CD31^−^) and positive surface markers (CXCR4^+^, β1-integrin^+^) was used to isolate MuSCs^13,14^ and genome-wide chromatin interactions were profiled through *in situ* Hi-C^15^ in biological duplicates (Figure 1A). The resulting proximity dataset replicates were processed with Juicer^16^, yielding highly reproducible contact maps (>0.98 Pearson correlation, Supp. Figure 1A-B). Pooled replicates comprised ~1.86×10^8^ chromosomal contacts for young MuSCs and ~1.56×10^8^ chromosomal contacts for aged MuSCs, of which ~63% were long-range (>20 kb) intra-chromosomal *cis*-interactions (Supp. Figure 1C). The Hi-C matrices exhibited sufficient sequencing depth to reveal chromatin structures such as topologically associated domains (TADs) and chromatin loops with up to 5 kb resolution under visual inspection in JuiceBox^17^ (Figure 1B). The contact maps were segmented into ~1 Mb regions associated with euchromatic (A) and repressive (B) chromatin compartments by analyzing the first eigenvector of the Pearson correlation matrices at 100 kb resolution^18^ (Supp. Figure 1D). This analysis partitioned the genome into ‘A’ and ‘B’ compartments in a 52.5/47.5 ratio in both young and aged MuSCs (Supp. Figure 1E) with similar compartmentalization strengths^19^ as measured by the average contact enrichment within and between compartments (Supp. Figure 1F). We observed minimal re-arrangement of A/B compartments, with <5% of the genome transitioning between compartments (Figures 1C-D). Consistent with previous findings, ‘A’ compartments displayed increased chromatin accessibility^5^ (ATAC-seq^20^, Figure 1E) and activating H3K4me3 signals^21^ compared to repressive ‘B’ compartments, and aged MuSCs displayed increased chromatin accessibility within static ‘A’ compartments (Supp. Figure 1G). RNA-seq datasets^5^ of age-matched MuSCs agreed with ‘A’ and ‘B’ chromatin compartment assignments, whereby genes within ‘A’ compartments showed increased expression relative to genes within ‘B’ compartments (2,059 differentially expressed genes in ‘A’; 154 in ‘B’, Figure 1F). In addition, gene expression within each compartment group displayed only subtle changes with aging apart from the static ‘A’ compartment, which showed a significant decrease in expression in aged MuSCs. Annotation of differentially expressed static ‘A’ genes revealed upregulation of transcripts in young MuSCs associated with cell-cycle checkpoints^22^ (*Cenpn, Ccne1, Cdc23*), SUMOylation of chromatin organizing proteins^23^ (*Pias1/2, Satb2, Rnf2*), and metabolism supportive of quiescence^24^, including fatty acid beta-oxidation (*Mecr, Hadha/b*, acyl-CoA dehydrogenase family) and the citric acid cycle (*Pdk1/2, Ldha, Pdha1*). In contrast, aged MuSCs showed increased expression in electron transport chain activity^24^ (NADH:ubiquinone oxidoreductase family) and response to interferon-beta (*Ifnar2, Igtp, Irf1*) (Supp. Figure 1H). Differentially expressed genes in the static ‘B’ compartment showed upregulation of G-protein coupled receptor (GPCR) activity (*Plppr1, Npy, Adra1b*) in aged MuSCs, which has been linked to stem cell fate regulation^25^, while genes expressed in young MuSCs were related to Rho GTPase activity^26,27^ (*Arhgap28/44, Pik3c3, Fgd4*) and cell migration (*Cdh13, Actc1, Vegfc*). Summing these results shows minimal plasticity in global chromatin architecture during MuSC aging, suggesting that changes in MuSC expression with old age may be conferred through altered local interactions in open chromatin compartments.

**Figure 1.**
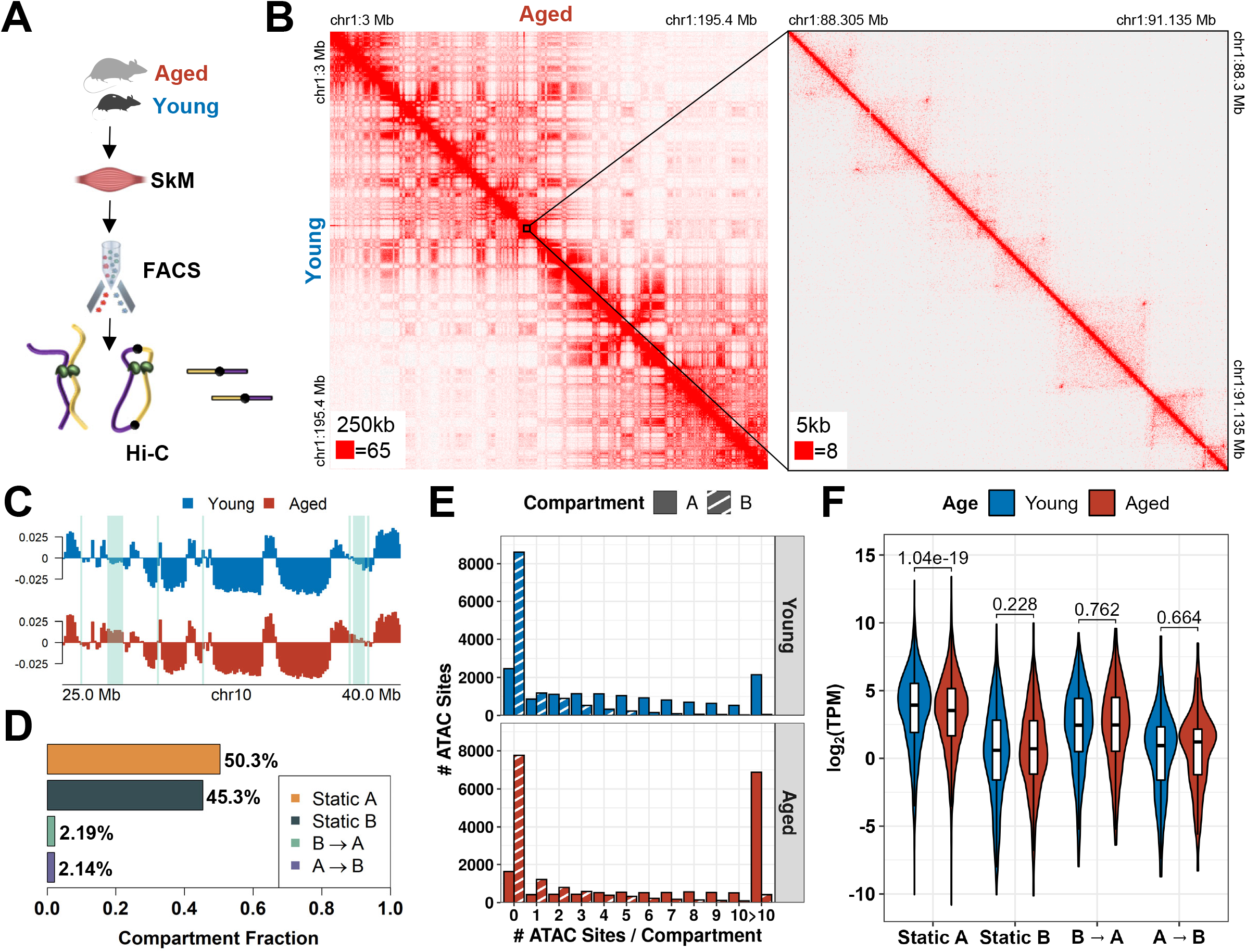
Global changes in 3D genome organization are largely static during muscle stem cell aging. **A)** Schematic of experiment. **B)** Normalized contact heatmaps of Young (lower triangle) versus Aged (upper triangle) MuSCs for chr1 at 250 kb resolution. A zoomed-in region is shown at 5 kb resolution. The maximum color map value for pixels is shown in the bottom left corner of each heatmap. **C)** Representative A/B compartment signal showing changes between young and aged MuSCs at 100 kb resolution. Compartment switches are highlighted in green. **D)** Quantification of 100 kb bins that switch compartments (Young to Aged). **E)** Distribution of ATAC-seq peaks in A and B compartments. **F)** Gene expression in log_2_(TPM) per A/B compartment in young and aged MuSCs.

### Local Chromatin Architecture is Altered During Muscle Stem Cell Aging

To investigate local changes in chromatin topology, we characterized TADs in 40 kb-resolution normalized contact matrices using HiCExplorer^28^. We identified 2,824 and 2,709 TADs in young and aged MuSCs, respectively (Figure 2A, Supp. Figure 2A). Integration of ATAC-seq and H3K4me3 signals revealed enrichments within TAD domains in an ‘A’ compartment-dependent manner (Figure 2B-C, Supp. Figure 2B). Consistent with previous reports^29^, TAD boundaries were enriched for CTCF motifs (Supp. Figure 2C), gene promoters, and transcription termination sites (Supp. Figures 2D-E), and expression of housekeeping genes was enriched at stable TAD boundaries (Supp. Figure 2F). TAD boundaries that were lost or gained with age were observed to rarely switch compartments from A→B or B→A (Figure 2D), and displayed reductions in both insulation strength (i.e. increased TAD separation scores) (Supp. Figures 2G-H) and gene expression (Supp. Figure 2I) relative to stable boundaries. TADs showed extensive restructuring with age, and re-arrangements were classified into shifts, splits, merges, and indeterminate rearrangements comprising some combination of the previous three classifications (Figures 2C,E-F). While these rearrangements showed no clear association with changes in gene expression within TAD domains or at TAD boundaries, we observed that boundaries that were gained with age (unique in aged MuSCs) were repositioned further from all nearby gene promoters compared to stable boundaries (Fisher’s test p-value < 2.2e-16, Supp. Figure 2J). Stable boundaries and those that were lost with age (unique in young MuSCs) were stationed at similar distances to gene promoters, indicating that TADs are potentially restructured during aging to create more space within TAD domains for regulatory interactions.

**Figure 2.**
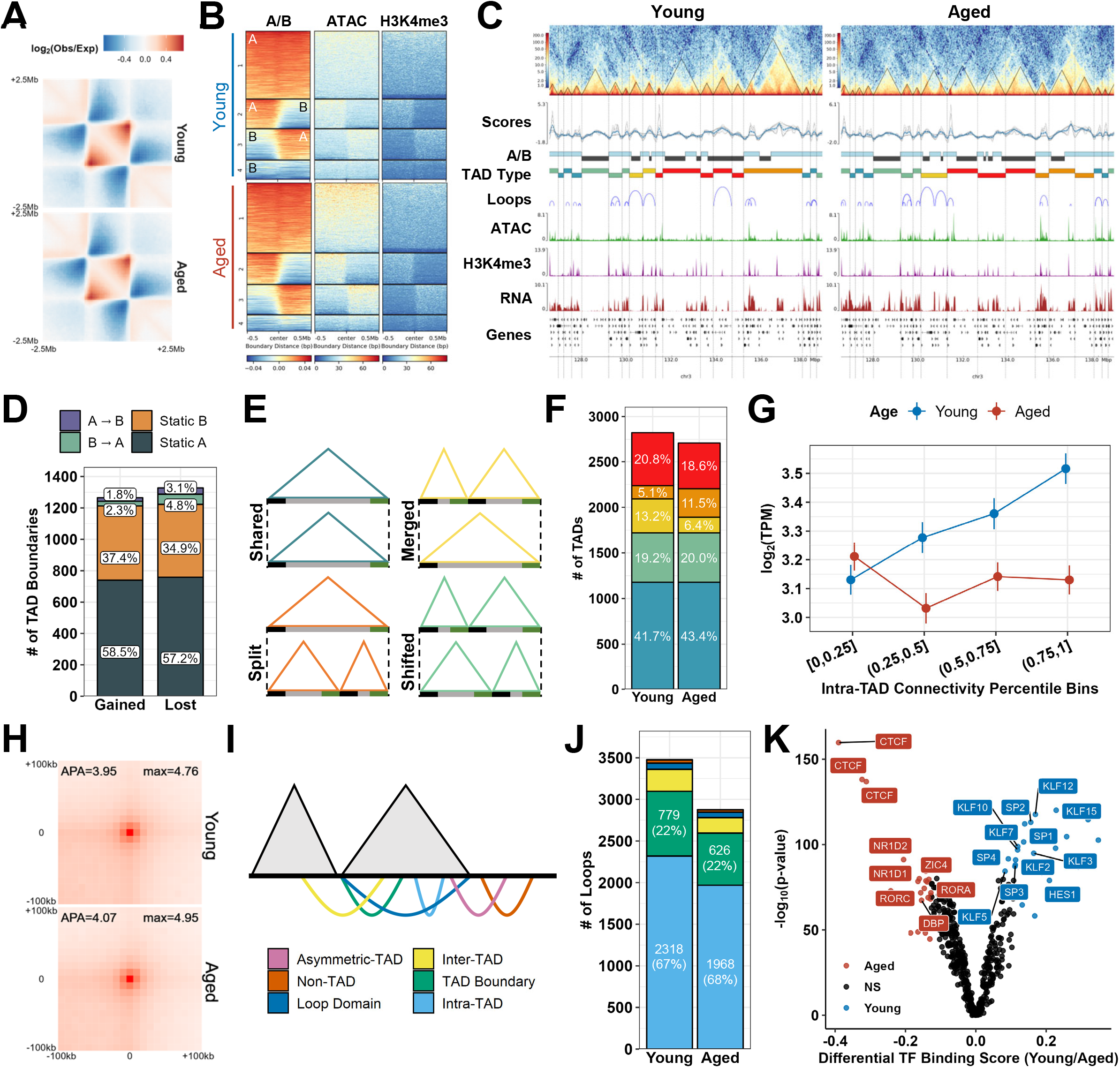
Muscle stem cells exhibit age-dependent changes in Topologically Associated Domains (TADs) and chromatin loops. **A)** Aggregate contact heatmaps at 40 kb resolution over a 5 Mb region centered on contact domains (TADs). **B)** Distributions of ATAC-seq and H3K4me3 signals (RPKM) across a 1 Mb region centered on TAD boundaries. K-means clustering (4 clusters) of the A/B compartment signal classified the boundaries by whether they fell within A/B compartments or divided compartment switching regions. **C)** Representative contact heatmaps log1p(observed/expected), TAD separation score tracks, A/B compartments (A, blue; B, gray), TAD domains, TAD boundaries (vertical lines and triangles), chromatin loops, ATAC-seq and H3K4me3 fold-change signal tracks, and gene expression in RPKM for young and aged MuSCs. TAD domains are colored by TAD rearrangement type. **D)** Distribution of TAD boundaries gained in aged and lost in young across A/B compartment groups. **E)** Cartoon showing definitions of rearranged TAD types in young MuSCs (top) relative to aged MuSCs (bottom). **F)** Proportions of TAD rearrangements. **G)** Gene expression (mean +/- SEM) in log_2_(TPM) against binned intra-TAD connectivity percentiles. **H)** Aggregate Peak Analysis (APA) of chromatin loops. **I)** Diagram and **J)** quantitation of chromatin loops classifications. **K)** Volcano plot of inferred binding activity of expressed (TPM>1) transcription factors (TFs) in sites of accessible chromatin at loop anchors. The top 5% differentially bound TFs are colored in red (aged) and blue (young).

To further understand age-dependent changes within TADs, we quantified the degree to which each TAD associates with itself by calculating intra-TAD connectivity, the mean number of reads per intra-TAD interaction as a fraction of reads for all *cis*-interactions with flanking TADs^11,30^ (Supp. Figure 2K). Intra-TAD connectivity complements TAD separation scores by assessing TAD compartmentalization rather than inter-TAD insulation. Intra-TAD connectivity was enriched in open ‘A’ compartments (Supp. Figure 2L), was positively correlated with activating chromatin features in both young and aged MuSCs (Supp. Figure 2M), and increased with age (Supp. Figure 2N). Increased intra-TAD connectivity was also strongly associated with increased gene expression in young MuSCs, but showed decreased association with age (Figure 2G). For example, genes related to cell cycle regulation (*Anapc1/4, Cdc25a, Mcm6, Prim2*) and lipid metabolism (*Acly, Ipmk, Sgms2, Aasdh, Pik3c3, Pi4k2b*) were encompassed by TADs with strong intra-TAD connectivity (top 20%) in both young and aged MuSCs but showed significantly upregulated gene expression in young MuSCs compared to aged. These results further suggest that in aged MuSCs, TADs self-interact in a stronger manner and alter TAD boundaries^31^ to facilitate increased interactions.

### Chromatin Loops Form Differential Gene Regulatory Units With Age

The changes within TADs during aging suggest that interactions between gene promoters and their cognate enhancers may be altered in aged MuSCs. To further explore contact domains between genes and their regulatory elements, we called chromatin loops using Hi-C Computational Unbiased Peak Search (HiCCUPS)^15^. We identified 3,478 and 2,877 chromatin loops in young and aged MuSCs (Supp. Figure 3A), respectively. High-scoring aggregate peak analysis (APA) plots confirmed the accuracy of the loop calls and revealed similar aggregate contact strengths across age (Figure 2H). ~90% of chromatin looping was constrained within individual TADs across aging (Figure 2I-J) and annotation of the loop anchors revealed enrichments for promoters, distal regulatory elements, and CTCF motifs (Supp. Figure 3B-D), with distal regulatory elements frequently connected at one anchor with a promoter or another regulatory element at the other (Supp. Figure 3E). Additionally, H3K4me3 levels and chromatin accessibility at promoters were enhanced when connected to distal regulatory elements enriched with the same chromatin features (Supp. Figure 3F). Gene expression within loop domains was also upregulated in young and aged MuSCs (Supp. Figure 3G), in line with previous observations that chromatin looping within TADs forms insulated neighborhoods that protect gene expression patterns from aberrant promoter-enhancer contacts that may form due to heightened intra-TAD connectivity^32^. To characterize the factors underlying expression changes associated with looping, we inferred differential binding of expressed transcription factors (TFs) at accessible sites within loop anchors using TF footprinting^33^ (Figure 2K, Supp. Figure 3H). We observed that young loop anchors were enriched for Notch-related TFs (*Hes1*) and multiple members of the Krüppel-like factors (*Klf*) and Specificity Factors (*Sp*) families, which have been shown to contribute to quiescence, chromatin organization, and regulation of proliferation^34–37^. In contrast, aged loop anchors showed enrichments for *CTCF, Zic4*, and TFs that regulate the circadian clock (*Nr1d1/2, Rora/c, Dbp*), which synchronizes pathways that are critical for MuSC homeostasis such as autophagy and responses to cell stress^38^, and have been recently shown to modulate chromatin topology^39,40^. Summing these results suggest aged MuSCs lose chromatin loops and TF binding that are associated with pathways that participate in maintenance of quiescence and cellular stress.

**Figure 3.**
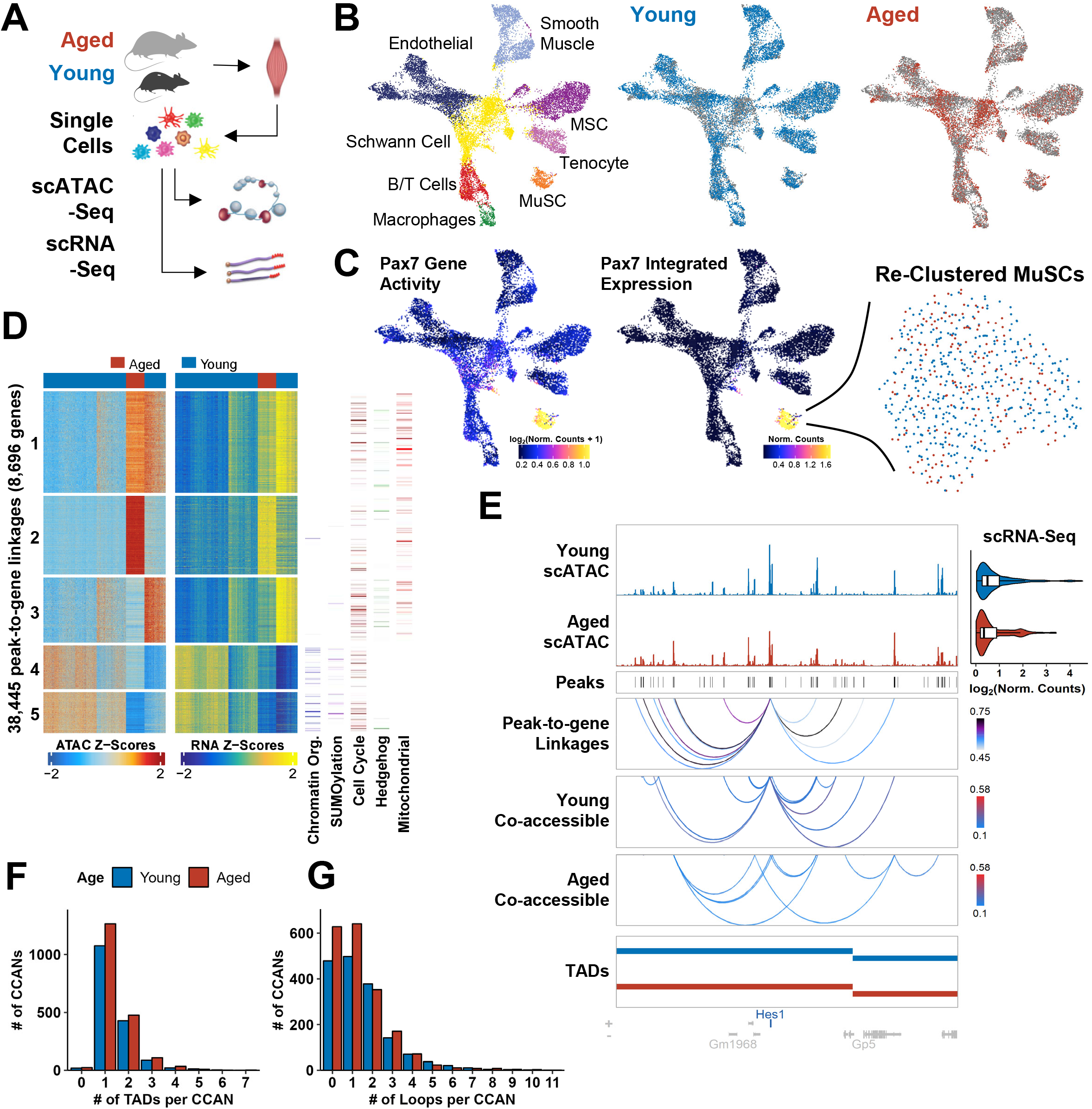
Single-cell multi-omic analysis of gene regulatory dynamics during muscle stem cell aging. **A)** Schematic of single-cell ATAC-seq (scATAC-seq) dataset generation and integration with single-cell RNA-seq (scRNA-seq) datasets. **B)** scATAC UMAP embeddings colored by cell type and age. **C)** scATAC UMAP embeddings colored by Pax7 gene activity scores and linked gene expression from integrated scRNA datasets. The identified MuSC cluster was reclustered and colored by age. **D)** Row-scaled heatmaps of statistically significant peak-to-gene linkages. Each row represents either chromatin accessibility at a distal site (left) or expression of the target gene (right). The columns represent cell aggregates colored by age. K-means clustering (5 clusters) reveals distinct regulatory networks between young and aged MuSCs. Heatmaps of representative Reactome pathways enriched in each cluster are shown to the right (see Methods). scATAC signal tracks, peaks, and peak-to-gene linkages (P2G) for *Hes1* colored by correlation scores. Cicero co-accessibility scores for the distal sites involved in these linkages are shown below colored by co-accessibility scores. Expression from integrated single cell RNA datasets are shown as violin plots. **F) and G)** Histograms of the number of CCANs constrained within individual TADs and **G)** loops.

### Single Muscle Stem Cell Multi-Omic Sequencing Shows Variation in Chromatin Hubs During Aging

Recent evidence suggests that cohesin-based extrusion of chromatin loops is short-lived^41^ and significant cellular heterogeneity exists at the level of promoter-enhancer contacts^42^. To increase the resolution of our analysis and further understand patterns of co-accessible sites within TADs, we performed single-cell ATAC-seq^43^ on mononucleated cells isolated from young (3 months) and aged (28-29 months) hind limb muscles (Figure 3A). We collected 15,263 cells (10,705 in young and 4,558 in aged) that passed quality control and filtering thresholds (Supp. Table 1, Supp. Figure 4A-C). Using a matrix of contiguous genomic tiles, we projected the datasets into low-dimensional space by iterative latent semantic indexing^44^, integrated them using Harmony^45^, and clustered the cells using the ArchR package^46^. We annotated each cluster by calculating gene-activity scores^47^, a metric derived from chromatin accessibility proximal to genes, and comparing scores at cell type marker genes with integrated single-cell transcriptomes from young and aged MuSCs that we previously generated^48^ and age-matched Tabula Muris Senis^49^ datasets (Supp. Figure 4D-G). This joint marker-based identification approach revealed similar cell types observed in other single-cell skeletal muscle atlases^50,51^ with differentially accessible sites in each cell type (Figure 3B, Supp. Figure 4H). For example, cells in the MuSC cluster were strongly enriched for gene activity scores and integrated gene expression for *Pax7* and *MyoD* compared to other cell types (Figures 3C, Supp. Figure 4D,G).

To identify putative distal regulatory elements, we called peaks in each cell type cluster using MACS2^52^ and iteratively created a union set of 48,842 peaks in ArchR. Most peaks lay within promoters (46%), with 21% and 24% laying in distal and intronic regions, respectively (Supp. Figure 5A). The MuSCs were then separately clustered (Supp. Table 1, Supp. Figure 4B-C) and MuSC-specific gene regulatory networks were identified by linking variation in chromatin accessibility at gene-distal sites with differences in local gene expression. This analysis revealed 38,445 significant peak-to-gene linkages with positive regulatory effects representing potential enhancer-gene interactions involving 14,240 peaks and 8,696 genes across young and aged MuSCs (FDR<1e-4, correlation>0.45) (Figure 3D). Clustering these linkages via k-means clustering revealed distinct cis-regulatory interactions between young and aged MuSCs with varying peak annotations (Supp. Figure 5A). Clustered linkages regulated molecular processes that we previously identified in the static ‘A’ chromatin compartment. For example, clusters 1 and 2 contained linkages enriched in aged MuSCs that regulated mitochondrial translation (Mitochondrial ribosomal protein families), respiratory electron transport (mitochondrial ATP synthase and NADH:ubiquinone oxidoreductase families), and DNA repair, while clusters 4 and 5 were dominated by young MuSCs and showed strong enrichment for genes that participate in chromatin organization (*Max* and *Myc*^53^, Histone lysine demethylase family), regulation of TP53 activity (*Mdm2, Atm, Trp53ka/b*), and SUMOylation of proteins (Figure 3D). All clusters contained linkages that regulate cell cycle checkpoints, and clusters 1 and 3 contained regulatory networks that both promoted (*Dhh, Smo, Gas1*) and inhibited (*Ptch1, Gli2, Gsk3*) Hedgehog signaling, which has been recently shown to support the regenerative capacity of MuSCs and is attenuated with aging^54^. Additionally, all linkages were predominantly confined to an individual TAD, indicating that these positive regulatory interactions between distal elements and their target genes are spatially restricted (Figure 3E, Supp. Figure 5B).

Next, we investigated whether transcriptional differences between young and aged MuSCs could be explained by differences in connectivity with cis-regulatory networks. We identified patterns of *cis*-co-accessibility between distal regulatory elements and promoters using Cicero^47^, yielding 27,015 and 31,512 pairs of co-accessible sites in young and aged MuSCs, respectively (Supp. Figure 5C). Consistent with previous reports^47,55,56^, co-accessible sites were more likely to be linked within the same TAD across co-accessibility thresholds compared to distance-matched peaks in separate TADs (Supp. Figure 5D). Similarly, we observed that >62% of sites within the average peak-to-gene linkage were co-accessible with each other, providing additional evidence that the linkages are part of spatially distinct networks. For example, linkages regulating differentially expressed genes, including *Hes1*, a downstream target of Notch signaling that was significantly upregulated in young MuSCs and differentially marked young chromatin loop anchors, showed differences in co-accessibility patterns, highlighting how variability in cis-regulatory networks can drive differential gene expression (Figure 3E).

To identify co-regulated chromatin hubs, we clustered the co-accessible sites into *cis*-co-accessibility networks (CCANs), yielding 1,648 and 1,922 CCANs in young and aged MuSCs, respectively. CCANs were primarily constrained within individual loops (30.3% in young, 33.2% in aged) and TADs (65.2% in young, 65.7% in aged, Figures 3F-G). We next applied a maximum weighted bipartite matching algorithm^47^ that identified 1,067 pairs of stable CCANs during aging. These CCANs accounted for >82% of all co-accessible sites in each dataset (Supp. Figure 5E) and encompassed >50% of all differentially expressed genes. Furthermore, we observed that the average matched CCAN shared only 36.9% of its constituent sites between young and aged MuSCs (Supp. Figure 5F) and that differential gene expression within these networks was significantly upregulated in young (Supp. Figure 5G). These results indicate that age-dependent expression changes are driven by co-accessible sites leaving or joining chromatin hubs. For example, the matched CCANs containing *Mta2*, a core member of the nucleosome remodeling and deacetylase (NuRD) complex^57^ that was enriched in young peak-to-gene linkages, and *Ndufc1*, a member of the NADH:ubiquinone oxidoreductase family upregulated in aged MuSCs, revealed altered connectivity with each gene body (Supp. Figure H-I). Together, these findings reveal extensive rewiring of connections between promoters and distal regulatory elements within chromatin hubs during aging that produce changes in gene expression. Changes in these interaction networks occurred primarily within individual TADs, which concentrate chromatin features that are supportive of gene expression, and chromatin loops, which insulate interactions and are associated with distinct TF networks.

## DISCUSSION

Aging encompasses declines in the functionality of multiple pathways^58,59^, yet how these different systems converge to modify genome organization in tissue resident stem cells has not been explored. To address this knowledge gap, we used a multi-omic approach to evaluate each level of chromatin architecture in young and aged MuSCs. We first gleaned how global genome structure is largely static in old age, and >95% of A/B compartments are invariant between young and aged MuSCs. These results are consistent with hematopoietic and neural development^60,61^, whereby stem cells display strong differences in phenotype but highly similar chromosomal compartments. In contrast, multiple types of TAD re-arrangements, increased inter-TAD interactions, and altered enhancer-promoter contacts within open TADs were observed in aged MuSCs. Many of these observed alterations displayed minimal impacts on gene expression, which is in line with previous studies that showed global loss of TAD boundaries through CTCF and/or cohesin deletion produced only mild perturbations to global gene expression^62–65^. Given that different systems such as in-vitro stem cell differentiation^11^ and acquisition of fibroblast senescence^66,67^ also show similar principles of stable global chromatin architecture, our results suggest that local alterations in genome folding within TADs are the primary drivers of changes in MuSC gene expression and associated age-related dysfunction.

Chromatin looping plays a key role in regulation of gene expression and occurs through a cohesin-based extrusion process that is stalled by DNA-bound CTCF proteins positioned in a ‘convergent’ orientation^15,41^. We detected a decrease in the number of chromatin loops in aged MuSCs and analysis of TF motifs at loop anchors revealed a loss in Notch-related and Krüppel-like TFs^68^ that are associated with MuSC quiescence and chromatin stability^34–37^. Notch has previously been demonstrated to regulate targets by repositioning enhancers^69^ and promoting increased interactions in enhancer ‘cliques’, which is consistent with our results in which cell cycle inhibitors and quiescence-related genes display enhanced contacts. The loss of these TF interactions in old age and enriched CTCF binding suggest that aging may promote alterations in the formation and stability of extruded loops. While a greater understanding of loop stability is needed, our results suggest MuSCs in old age may prematurely activate to generate ATP^70^ to maintain or recreate lost loop domains. As a consequence of loop dysregulation in old age, interactions between promoters and their concomitant regulatory elements are modified, driving alterations in TF binding and the affinity of distal regulatory elements for their target genes and associated expression levels.

Developmental maturation of stem cells has been shown to be driven by the combinatorial action of multiple enhancers, resulting in increased enhancer–promoter interactions of specific genes^61^ and variations in TF binding^71^. Our results are consistent with this stem cell continuum, showing that old age results in alterations of promoter-enhancer elements with accessible sites frequently leaving or joining chromatin hubs that maintain chromatin organization and quiescence. Our findings show that rewiring of chromatin within TADs and loops underpins aberrant gene expression programs associated with pathological aging. Future work in this area will resolve additional structural features such as TAD stripes^70^ and interactions with other nuclear structures such as speckles and nucleoli.

## ACKNOWLEDGEMENTS

The authors thank Arima Genomics for assistance with Hi-C, and the University of Michigan DNA Sequencing Core for assistance with sequencing. The authors also thank Anna Shcherbina, and Jie Liu for insights into bioinformatics analysis, and members of the Parker and Aguilar laboratories. Research reported in this publication was supported by a National Science Foundation CAREER award (2045977), partially supported by the National Institute of Arthritis and Musculoskeletal and Skin Diseases of the National Institutes of Health under Award Number P30 AR069620 (CAA), the 3M Foundation (CAA), American Federation for Aging Research Grant for Junior Faculty (CAA), the University of Michigan Geriatrics Center and National Institute of Aging under award number P30 AG024824 (CAA), and the National Institute for Biomedical Imaging and Bioengineering Training Award T32 EB005582 (BAY). The content is solely the responsibility of the authors and does not necessarily represent the official views of the National Institutes of Health.

## Author contributions

J.L., K.M.S., and P.M.F. performed experiments. B.A.Y. analyzed data. C.A.A. designed the experiments. S.C.J.P. contributed reagents / tools. B.A.Y., and C.A.A. wrote the manuscript.

## Materials & Methods

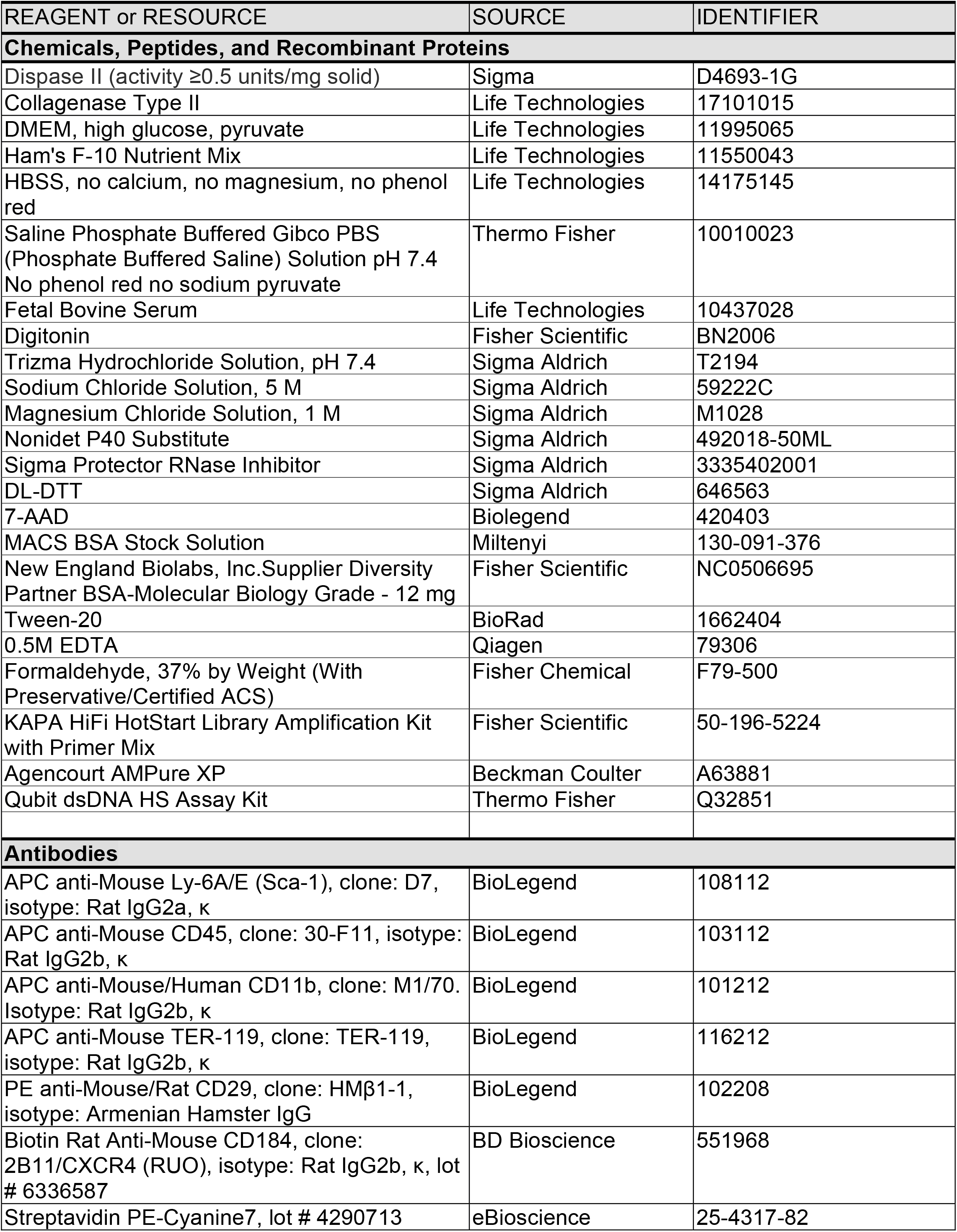
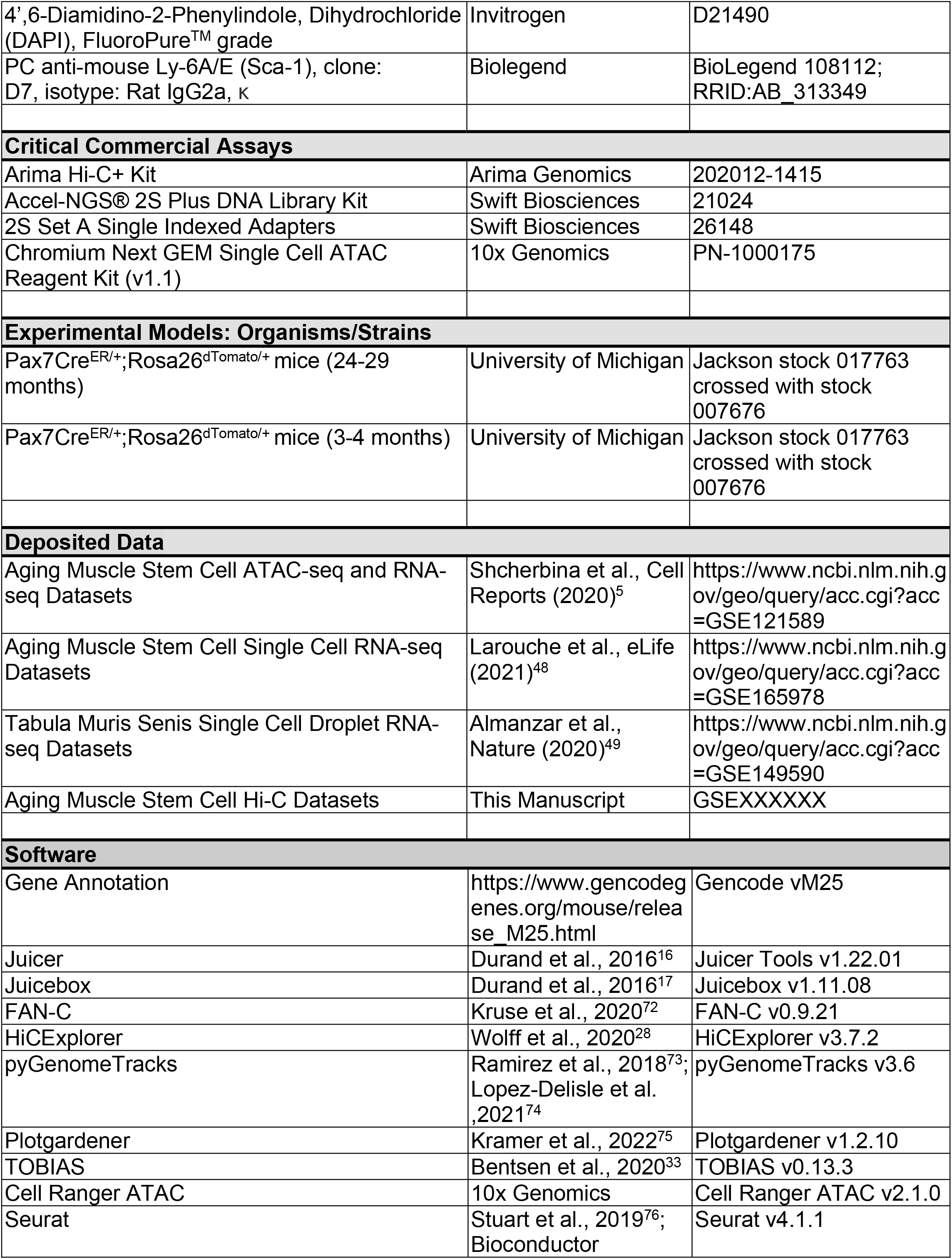
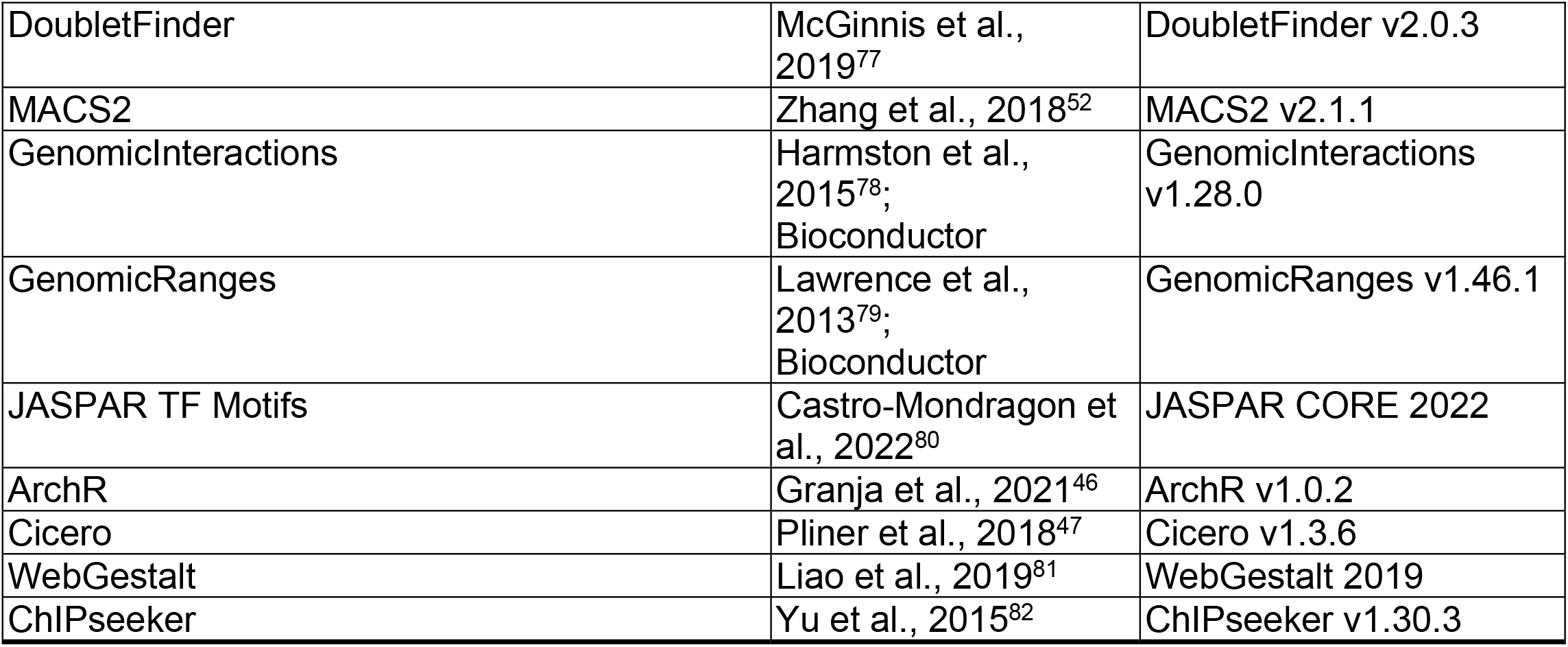

### Hi-C Library Preparation and Sequencing

#### Animals

Young (4 months) and aged (24-26 months) Pax7Cre^ER/+^;Rosa26^TdTomato/+^ female mice were obtained from a breeding colony at the University of Michigan (UM). All mice were fed normal chow ad libitum and housed on a 12:12 hr light-dark cycle under UM veterinary staff supervision. All procedures were approved by the University Committee on the Use and Care of Animals at UM and were in accordance with the U.S. National Institute of Health (NIH).

#### Isolation and crosslinking of muscle stem cells

MuSCs were isolated by FACS as previously described^5^. In brief, hind limb muscles were harvested from mice using sterile surgical tools. Muscle tissue was minced and digested in 20 mL of a digestion solution (2.5 U/mL Dispase II and 0.2% [~5500 U/mL] Collagenase Type II in Dulbecco’s modified Eagle medium [DMEM]). Samples were incubated at 37°C for 60 min. Once the digestion was complete, 20 mL of F10 media containing 20% heat inactivated FBS was added into each sample to inactivate enzyme activity. The solution was then filtered through a 70 µm cell strainer into a new 50 mL conical tube and centrifuged at 350× *g* for 5 min. Subsequently, the protocol for crosslinking low input mammalian cells was followed, as written in the Arima-HiC kit workflow provided by Arima Genomics Inc (San Diego, CA). Briefly, the pellets were re-suspended in 1X PBS and 37% formaldehyde was added to obtain a final concentration of 2% formaldehyde. The samples were inverted and incubated at RT for 10 mins. Arima-Hi-C Stop Solution 1 was added to samples, and they incubated for an additional 5 mins. The samples were then placed on ice to incubate for 15 mins. Cells were pelleted by centrifugation for 5 min at 500 × *g*, the supernatant was discarded, and cells were resuspended in staining media (2% heat-inactivated FBS, 2 mM EDTA in Hank’s buffered salt solution) and antibody cocktail containing Sca-1:APC (1:400), CD45:APC (1:400), CD11b:APC (1:400), Ter119:APC (1:400), CD29/β1-integrin:PE (1:200), and CD184/CXCR-4: BIOTIN (1:100) and incubated for 30 min on ice in the dark. Cells and antibodies were diluted in 3 mL of staining solution, centrifuged at 350× *g* for 5 min, and supernatants discarded. Pellets were re-suspended in 200 µL staining solution containing PECy7:STREPTAVIDIN (1:100) and incubated on ice for 20 min in the dark. Again, samples were diluted in 3 mL staining solution, centrifuged, supernatants discarded, and pellets re-suspended in 200 μL staining buffer. Samples were filtered through 70 μm cell strainers before the FACS.

#### Preparation of Hi-C Libraries & Sequencing

In situ Hi-C was performed using the Arima-HiC Protocol. Approximately 75,000-100,000 sorted muscle cells were crosslinked, permeabilized and chromatin digested as specified by the manufacturer in biological replicates. Restriction fragment ends were then labeled with biotinylated nucleotides and proximal DNA ligated, followed by reversal of cross-links. The DNA was then sheared using Diagenode’s Bioruptor sonicator for 4 cycles of 15 sec on, 90 sec off, then briefly centrifuged. An additional 4 cycles of sonication were performed after centrifugation. The resultant DNA was size-selected using AMPure XP beads followed by verification with the Arima Quality Control 2 assay. Biotinylated junctions were then magnetically isolated with streptavidin coated beads, followed by end-repair, A-tailing, sequencing adaptor ligation and PCR amplification. The resultant libraries were then size-selected using AMPure XP beads and sequenced with 150 bp paired-end reads at a depth of 750 million reads per library on an Illumina NovaSeq flow cell.

### Hi-C Data Preprocessing

Sequence quality metrics were checked in all samples using FastQC (v0.11.9, https://www.bioinformatics.babraham.ac.uk/projects/fastqc/) before downstream processing. Read adapters were trimmed using Cutadapt^83^ (v2.6). Only trimmed reads >20bp were retained and 25 bases were trimmed from the 3’ ends. Paired-end reads for each replicate were then split and each mate was aligned as single-end reads to the mouse reference genome (mm10) using BWA aln and samse (v0.7.17-r1188). Uniquely mapped reads classified as “optimal” or “suboptimal” by BWA with a maximum edit distance of 3 and MAPQ>=30 were retained. Paired mates were then re-joined and duplicate reads removed. Each chromosome was split into 5kb bins and the number of interactions in each bin was counted. Interactions were eliminated if they occurred on the same fragment (i.e. overlapping reads on the same chromosome) or if the distance between the start coordinates of each read pair was <1kb. All remaining reads were kept if they participated in at least one interaction. The resulting paired contact maps were converted to .hic files using the pre command in Juicer Tools^16^ (v1.22.01) with a file of Arima restriction sites created with the generate_site_positions.py script. The .hic file was created with the following 9 base-pair-delimited resolutions: 2.5Mb, 1Mb, 500kb, 250kb, 100kb, 50kb, 40kb, 25kb, 10kb, and 5kb.

### Hi-C Quality Control

Pairwise Pearson correlations were computed between Knight-Ruiz-normalized replicates at 250kb-resolution with HiCExplorer^28^ (v3.7.2) after converting .hic files to .cool files using the hicConvertFormat command. All replicates showed Pearson correlations <0.98 and replicates were pooled for downstream processing. The fraction of inter-chromosomal interactions in pooled replicates was ~23% while ~63% of interactions occurred more than 20kb apart and ~14% of interactions occurred less then 20kb apart, indicating good quality libraries. Replicates also showed the same distributions of count enrichment at different genomic ranges as shown by the hicPlotDistVsCounts command with a maximum distance from the diagonal of 30 Mb.

### A/B Compartment analysis

A/B compartments were identified from the first eigenvectors of the Pearson correlation matrices of 100kb-resolution contact matrices using the ‘compartments’ command in the FAN-C^72^ (v0.9.21) package. The sign of the first eigenvector was oriented by the average GC content of each domain such that domains with higher GC content were assigned positive signs (A compartment) while domains with lower GC content were assigned negative signs (B compartment). The compartment eigenvector BED files were then exported from FAN-C and analyzed in R to identify compartments that were static or shifted between young and aged MuSCs. Saddle plots were generated using as an additional output of the ‘compartments’ command.

### TAD Calling and Characterization

#### Identification of TADs

TADs were called from Knight-Ruiz-normalized contact matrices at several resolutions (10kb, 40kb, 100kb, 250kb, and 500kb) and FDR thresholds (0.1, 0.05, 0.01, 0.005, and 0.001) using the ‘hicFindTADs’ command in the HiCExplorer package with default parameters. The final set of TADs and associated boundaries (40kb resolution, FDR<0.01) was selected by visual inspection with pyGenomeTracks^73,74^ (v3.6) and comparisons of the size and number of TADs at each pair of parameters. Aggregate contact matrices around TAD domains were plotted using FAN-C aggregate.

#### Calculation of Intra-TAD Connectivity

Intra-TAD connectivity was calculated as the mean number of reads per intra-TAD interaction as a fraction of reads for all *cis*-interactions with flanking TADs. The number of contacts within TADs and between TADs was collected using the ‘hicInterIntraTAD’ command in HiCExplorer.

### Chromatin Loop Analysis

#### Loop Calling

Chromatin loops were identified using the HiCCUPS algorithm^15^ with default parameters for medium resolution maps (hiccups -m 512 -c (all chromosomes) -r 5000,10000,25000 -k KR -f .1,.1,.1 -p 4,2,1 -i 7,5,3 -t 0.02,1.5,1.75,2 -d 20000,20000,50000 /path/to/hic/file path/to/loops/files). The algorithm was run in a Google Colaboratory notebook with a GPU hardware accelerator. The merged loop list across all resolutions (i.e. 5kb, 10kb, 25kb) were used for downstream analyses.

#### Aggregate Peak Analysis (APA)

Aggregate peak analysis^15^ was used with default settings to evaluate focal contact enrichment at merged loops. We summarized the genome-wide APA analysis using the ratio of the central pixel to the mean of the mean of the pixels in the lower left corner. For visualization, we used the genome-wide normalized APA matrices at 10kb resolution. In these matrices, each submatrix was normalized by its mean such that the mean of the submatrix was 1.

### Feature Annotation and Gene Set Enrichment Analysis

TAD boundaries and loop anchors were annotated for genomic features using ChIPseeker (v1.30.3)^82^ with a promoter region of +/-1kb. Overlapping genomic annotations were resolved in order of decreasing priority as follows: Promoter, 5UTR, 3UTR, Exon, Intron, Downstream, Intergenic. Gene ontology (GO) and Reactome term enrichments were performed using over-representation (ORA) and gene set enrichment (GSEA) analysis^84^ using WebGestalt 2019^81^. Tested gene sets were limited to those containing between 5 and 2000 genes. Significant term enrichment thresholds were set at FDR < 0.05. For GSEA, genes were ranked according to the signal to noise metric^84^.

### CTCF Motif Finding

CTCF motifs were identified across the mm10 genome using scanMotifGenomeWide.pl from the HOMER suite^85^ with the following HOMER position weight matrix (PWM) for CTCF: >ANAGTGCCACCTGGTGGCCA CTCF(Zf)/CD4+-CTCF-ChIP-Seq(Barski_et_al.)/Homer,BestGuess:CTCF(Zf)/CD4+-CTCF-ChIP-Seq(Barski_et_al.)/Homer(1.000) 8.704837 -6.281855e+03 0 15000.0,4645.0,2877.2,2765.0,0.00e+00

0.447 0.221 0.181 0.151

0.037037037037037 0.34034034034034 0.236236236236236 0.386386386386386

0.499 0.061 0.33 0.11

0.033033033033033 0.377377377377377 0.528528528528528 0.0610610610610611

0.023023023023023 0.378378378378378 0.005005005005005 0.593593593593594

0.061 0.005 0.887 0.047

0.0790790790790791 0.905905905905906 0.005005005005005 0.01001001001001

0.002 0.994 0.001 0.003

0.501501501501502 0.475475475475475 0.00700700700700701 0.016016016016016

0.002 0.527 0.004 0.467

0.003 0.995 0.001 0.001

0.03 0.036 0.004 0.93

0.382 0.042 0.446 0.13

0.02 0.273 0.686 0.021

0.047 0.039 0.014 0.9

0.002 0.001 0.995 0.002

0.04 0.034 0.873 0.053

0.161161161161161 0.527527527527528 0.0610610610610611 0.25025025025025

0.277 0.428 0.117 0.178

0.541 0.092 0.267 0.1

CTCF motifs were enumerated in 5kb windows across loop domains and in 20kb windows across TAD domains. Loop and TAD domains were divided into 40 bins and the average number of motifs was plotted within each bin.

### Integration with RNA-Sequencing

#### Gene Expression Quantification

Paired-end RNA-seq data from MuSCs^5^ was aligned to the mm10 reference genome with the STAR algorithm (STAR_2.5.0a)^86^ using default parameters. RSEM^87^ quantification was applied to the aligned reads. The full set of flags used in the STAR command is as follows: STAR --genomeLoad NoSharedMemory --outFilterMultimapNmax 20 --alignSJoverhangMin 8 -- alignSJDBoverhangMin 1 --outFilterMismatchNmax 999 –outFilterMismatchNoverReadLmax --alignIntronMin 20 --alignIntronMax 1000000 --alignMatesGapMax 1000000 -- outSAMunmapped Within --outFilterType BySJout --outSAMattributes NH HI AS NM MD -- outSAMtype BAM SortedByCoordinate --quantMode TranscriptomeSAM --sjdbScore 1 -- limitBAMsortRAM 60000000000 --twopassMode Basic --twopass1readsN -1

#### Gene Expression Analysis

Gene expression analysis between young and aged samples was performed by limma^88^ analysis in R. The Expected Counts from RSEM were transformed to counts per million using the voom^89^ R package with a design formula: Count~Age, with Age={Young, Aged}. Surrogate variable analysis was performed with the SVA package^90^ using a null model of voom$E~ 1, and a design matrix of voom$E~Day+Age. Contributions from the surrogate variables were quantified and removed from the voom$E data matrix. Pairwise Pearson and Spearman correlation values were computed between all sva-corrected replicates. Any replicate that had r < 0.9 with other replicates for a given sample was excluded from further analysis. Expression analyses were performed on known genes filtered to those with TPM (transcript per million) values greater than 0 in at least one sample. Differentially expressed genes were identified as those with log_2_(fold-change)>1.5 and adjusted p-values<0.05. Compartments, loop anchors, and TADs were annotated with mm10 genes (GRCm38.p6 GENCODE M25) by identifying genes whose promoter regions (TSS +/- 1 kb) intersected each set of regions.

### Integration with ATAC-seq and H3K4me3 ChIP-seq Datasets

#### ATAC-seq Data Processing and Analysis

ATAC-seq datasets from young and aged MuSCs were collected previously^5^. The ATAC-seq samples were analyzed with the ENCODE ATAC-seq processing pipeline^91^ (https://github.com/ENCODE-DCC/atac-seq-pipeline, version 1.1.7). Read adapters were trimmed with the cutadapt algorithm^83^ and aligned to the mm10 reference genome using Bowtie2^92^. Duplicates were then marked using Picard (https://broadinstitute.github.io/picard/) and removed from the aligned reads using SAMtools^93^. The resulting BAM files were filtered to remove unmapped or unpaired reads and reads with MAPQ scores below 30. The MACS2^52^ peak caller was used to call peaks from the aligned ATAC-seq samples and the naive overlap peak set from all replicates for a given sample was used for downstream processing.

#### H3K4me3 ChIP-seq Data Processing and Analysis

FASTQ files for H3K4me3 ChIP-seq datasets from young and aged MuSCs were obtained from Liu et al., 2013^21^ and processed with the ENCODE ChIP-seq processing pipeline^91^ (https://github.com/ENCODE-DCC/chip-seq-pipeline2, version 2.1.1) using the “histone” option. Paired-end read adapters were trimmed using Trimmomatic^94^ and aligned to the mm10 reference genome using Bowtie2^92^. Duplicates were then marked with Picard (https://broadinstitute.github.io/picard/) and removed with SAMtools^93^. The resulting BAM files were filtered to remove unmapped or unpaired reads and reads with MAPQ scores below 30.

#### Signal Track Coverage

RPKM-normalized signal tracks were generated from merged ATAC-seq and H3K4me3 bam alignments using the ‘bamCoverage’ command in the deepTools suite (v.3.3.0)^95^ with a bin size of 1. Reads with the 780 SAM flag and mapping qualities <30 were excluded. The mean signals in chromatin compartment groups, TADs, and chromatin loop anchor were calculated using the ‘multiBigWigSummary’ command in deepTools. Blacklisted regions^96^ were excluded from these calculations.

#### Clustering of Signals Across TAD Boundaries By Compartment

The mean signal of the first eigenvector of the Hi-C Pearson correlation matrix was calculated in 10 kb-bins across 1 Mb regions flanking TAD boundaries using the ‘computeMatrix’ command in the deepTools suite. K-means clustering of the resulting matrices into 4 clusters using the ‘plotHeatmap’ command partitioned the TAD boundaries into those that lay within A/B compartments or spanned A/B compartment switches. Matrices generated from ATAC-seq and H3K4me3 fold-change signals in the same regions per cluster were plotted as heatmaps.

#### Pearson Correlation of ATAC-seq and H3K4me3 Signals with TADs

Mean fold-change signals from ATAC-seq and H3K4me3 data were calculated in 1 kb bins within TAD domains scaled to 15 kb regions and in flanking 15 kb regions using the computeMatrix command. Pearson correlation between the matrix columns for each signal was calculated in R, producing 45×45 matrices. Heatmaps were then generated for each correlation matrix.

#### Transcription Factor Footprint Analysis

TF motifs were downloaded from the JASPAR CORE 2022 database^80^ and filtered to those expressed (TPM≥1) in our bulk RNA-seq datasets. Heterodimer motifs were considered expressed if at least one partner was expressed.

All footprint analyses within chromatin loop anchors were performed using the TOBIAS toolkit (v0.13.3)^33^. The bam mm10 alignments from the ENCODE ATAC-seq pipeline were corrected for Tn5 enzymatic bias using the ‘ATACorrect’ function. Blacklisted regions^96^ were excluded from the bias estimations. To perform differential TF binding analysis, a union set of ATAC peaks was generated across young and aged MuSCs using the bedtools merge command. Continuous footprinting scores for the young and aged bias-corrected ATAC signals were then calculated across the union peak set using the ‘ScoreBigwig’ function. Estimates of TF binding positions were calculated using the ‘BINDetect’ function in the subset of union ATAC peaks that lay within merged loop anchors. Significant TFs were identified as those above the 95^th^ percentile or below the 5^th^ percentile of differential binding scores or those above the 95^th^ percentile of -log10(p-values).

### Generation and Processing of Skeletal Muscle Single Cell ATAC and RNA Datasets *Single Cell RNA Datasets*

Single cell RNA libraries (10x Genomics) from FACS-isolated MuSCs were previously generated from uninjured young (2-3 months) and aged (22-24 months) hindlimb muscles^48^. We downloaded a pre-processed droplet single-cell RNA dataset from limb muscle generated by the Tabula Muris Senis (TMS) atlas^49^ and converted it to a Seurat (v4.1.1)^97^ object. 3-month and 24-month data from the TMS object were used for downstream analyses.

TMS datasets were considered pre-processed and exempt from further quality control filtering. Our previously generated MuSC datasets were filtered to retain high-quality cells expressing >500 unique molecular identifiers (UMIs), between 300 and 4,000 genes, <10% mitochondrial reads, and gene complexities (log_10_(# genes) / log_10_(# UMIs)) > 0.8.

#### Integration of Single Cell RNA Datasets

All datasets were separately log-normalized and scaled to 10,000 using the ‘NormalizeData’ function, and feature selection was performed using the “vst” method in the ‘FindVariableFeatures’ function with the number of top variable features set to 2,000. Contributions from the number of UMIs per cell were regressed out. We additionally regressed out contributions from the percentage of mitochondrial reads in our previously generated datasets. We performed doublet analysis using DoubletFinder (v.2.0.3)^98^ with a 4.8% expected doublet rate and found no clear evidence of doublets in any dataset. The datasets were then integrated using the canonical correlation analysis (CCA) implementation in Seurat with default parameters.

The integrated dataset was re-scaled and principal component analysis (PCA) was performed on the 2,000 most variable genes. The number of PCs used to represent the dataset was quantitatively determined by identifying the minimum between the last PC where the change in variation was more than 0.1% and the last PC associated with <5% of variation that also accounted for >90% of the cumulative variation in the dataset (33 PCs). These PCs were used to compute the nearest neighbor graph (FindNeighbors) and were visualized using the ‘uwot’ implementation of Uniform Manifold Approximation and Projection (UMAP)^44^ with min.dist=0.5, n.neighbors=30, and the ‘cosine’ distance metric. Cells were clustered using the Louvain clustering algorithm implemented by Seurat’s ‘FindCluster’ function with a resolution of 0.1.

#### Single Cell RNA Cell Type Annotation

Differentially expressed genes between clusters in the integrated dataset were identified using the ‘FindAllMarkers’ function in Seurat with a Wilcoxon test to compare genes with >0.25 log_2_(fold-change) that were expressed in >10% of cells in each cluster. Clusters were annotated by cell type according to differentially expressed marker genes. Expression of marker genes was verified in each dataset within the integrated object using the ‘FindConservedMarkers’ function with the same parameters.

#### Skeletal Muscle Nuclei Extraction and Preparation of Single Cell ATAC Libraries

Young (3 months old) and aged (28-29 months old) Pax7Cre^ER/+^;Rosa26^TdTomato/+^ female mice were obtained from a breeding colony at UM. All mice were fed normal chow ad libitum and housed on a 12:12 hour light-dark cycle under UM veterinary staff supervision. All procedures were approved by the University Committee on the Use and Care of Animals at UM and the Institutional Animal Care and Committee and were in accordance with the U.S. National Institute of Health (NIH).

Mouse hindlimb muscles were extracted and placed into separate petri dishes containing ice-cold PBS. Using surgical scissors, muscle tissues were minced and placed into 20mL of digestion buffer (DMEM with Collagenase type II (0.2%) and Dispase II (2.5U/mL)) per mouse. Samples were placed on a shaker in a 37°C incubator for 1.5 hours and mixed by pipette every 30 minutes. The enzymes were then inactivated by addition of 20mL 20% heat-inactivated fetal bovine serum (HI-FBS) in Ham’s F10 media. The solution was passed through a 70um cell strainers, centrifuged, washed in PBS containing 3% BSA, and nuclei isolation was performed according to 10x Demonstrated Protocol CG000375 Revision B starting at step 1.1c. Nuclei from two age-matched mice were pooled prior to sorting, at step 1.1n, and 7-AAD+ nuclei were sorted on a Sony MA900 or MoFlo Astrios 3. Sorted nuclei were then permeabilized according to the same protocol. 10,000 permeabilized nuclei were loaded onto the 10x Genomics Chromium single cell controller and single nuclei were captured into nanoliter-scale gel bead-in-emulsions (GEMs). Single nuclei ATAC library constructions were performed using the ATAC-seq NextGEM kit (10x Genomics). All libraries were submitted for 51×51bp paired-end sequencing on a NovaSeq 6000 with 50,000 targeted reads per cell.

#### Single Cell ATAC Processing

Raw sequencing data were demultiplexed and converted to FASTQ files using the ‘bcl-convert’ command (Illumina, v3.9.3) and aligned to the mm10 reference genome (refdata-cellranger-arc-mm10-2020-A-2.0.0) using the ‘cellranger-atac count’ command (10x Genomics, cellranger-atac-2.1.0).

scATAC fragment files from all samples were processed simultaneously using the ArchR package (v1.0.2). Low-quality cells were removed based on transcription start site (TSS) enrichment (>4) and the minimum number of unique fragments per cell (>1,000). Doublets were filtered using the ‘addDoubletScores’ and ‘filterDoublets’ functions with default parameters (5.5% of total cells removed). Next, a cell-feature matrix of 500bp genome-wide tiles was used to create a low-dimensional representation of the dataset through an iterative latent semantic indexing (LSI) approach (‘addIterativeLSI’ function; 3 iterations; 30 LSI dimensions; 25,000 variable features; and 0.1 clustering resolutions). Batch effects were corrected using Harmony (‘addHarmony’ with default parameters)^45^ and LSI dimensions that were highly correlated with sequencing depth were excluded from downstream analyses (Pearson correlation>0.75). Cells were visualized using the ‘uwot’ implementation of UMAP^44^ embeddings using Harmony-corrected dimensions with the cosine metric, nNeighbors=30, and minDist=0.3 (‘addUMAP’), and cells were clustered using the FindClusters function in Seurat (v4.1.1)^76^ with resolution 0.4 (‘addClusters’).

#### Single Cell ATAC Cell Type Annotation and Imputed Gene Expression

Prior to labeling scATAC clusters with scRNA annotations, gene activity scores were inferred from scATAC data using the ‘addGeneScoreMatrix’ function in ArchR. These scores predict gene expression levels based on the accessibility of nearby regulatory elements. We identified marker genes for each scATAC cluster based on gene activity scores using a Wilcoxon test with the ‘getMarkerFeatures’ function. A suitable null background for this analysis was corrected for biases from TSS enrichment and the log_10_(# of unique fragments) per cell. Cell types were tentatively assigned to clusters based on these marker genes (FDR≤0.01, log_2_(fold-change)≥1.25), including MuSCs, which were marked by high Pax7 and Myod1 gene activity scores. To visualize these scores on UMAP embeddings, we smoothed gene activity scores across neighboring cells using MAGIC (Markov affinity-based graph imputation of cells) imputation^99^. We then used Seurat’s CCA implementation to assign the most similar scRNA cell to each scATAC cell by comparing the scATAC gene activity and scRNA gene expression matrices (‘addGeneIntegrationMatrix’), effectively transferring scRNA cell type labels to the scATAC cells. Gene expression profiles from scRNA cells were assigned to the corresponding scATAC cells. This integration was constrained to ensure that the putative scATAC MuSC cluster aligned with the scRNA MuSC cluster. scATAC clusters were then merged according to cell type annotation.

#### Single Cell ATAC Peak Calling

To overcome the sparsity of scATAC data, we generated pseudo-bulk replicates for each cell type cluster (‘addGroupCoverages’) and pseudo-bulk peak calling was performed using MACS2^52^ (‘addReproduciblePeakSet’). Peaks were merged into a union set in ArchR to create a single peak matrix (‘addPeakMatrix’) for the entire scATAC dataset. Differential peak accessibility (FDR≤0.1, log_2_(fold-change)≥1) across cell types was verified using the ‘getMarkerFeatures’ function with the same parameters as the analysis performed for gene activity scores. A heatmap of marker peaks was plotted using the ‘plotMarkerHeatmap’ function.

### Multi-omic Profiling of Muscle Stem Cells

#### Reclustering of MuSCs

The MuSC cluster of the scATAC dataset with linked scRNA expression was isolated and reclustered using the same iterative LSI procedure described previously (‘addIterativeLSI’) with 2 iterations and 500bp genome-wide tiles. Harmony batch correction was not necessary for these cells. Pseudo-bulk replicates were generated for young and aged MuSCs and a unified pseudo-bulk peak set was identified using MACS2^52^ and the ArchR merging procedure.

#### Identification and Clustering of Peak-to-Gene Linkages

Peak-to-gene correlation analysis was performed to identify putative gene-enhancer regulatory linkages from scATAC and scRNA data. This was accomplished with the ‘addPeak2GeneLinks’ function in ArchR with the first 30 LSI dimensions and a maximum distance between gene promoters and the center of accessible sites of 500kb. This function requires the generation of low-overlapping cell aggregates to overcome the sparsity of scATAC data. We slightly modified the cell aggregation step of ArchR’s implementation such that aggregates were composed solely of young or aged MuSCs. This modification allowed us to compute peak-to-gene correlations across the MuSC cluster while retaining age-specific contributions. The resulting linkages were filtered (correlation>0.45, FDR<1e-4) for downstream analyses. Linkages were clustered into 5 k-means clusters and visualized using the ‘plotPeak2GeneHeatmap’ function. Groups of linkages dominated by aged or young MuSCs could thus be visually identified.

#### Gene Set Annotation of Peak-to-Gene Linkages

Enriched GO and Reactome terms for linked genes in each cluster were analyzed using over-representation analysis in WebGestalt 2019^81^ with a FDR threshold of 0.1. Gene sets were limited to those containing between 5 and 2000 genes. Unique genes in representative enriched Reactome terms per cluster were aggregated and shown as heatmap annotations (Figure 3D). The selected terms were related to chromatin organization (R-MMU-4839726, R-MMU-3247509), SUMOylation of proteins (R-MMU-4570464, R-MMU-2990846), cell cycle regulation (R-MMU-69206, R-MMU-1640170, R-MMU-69278, R-MMU-69275, R-MMU-174184, R-MMU-69231, R-MMU-68882), Hedgehog signaling (R-MMU-5358351, R-MMU-5610787, R-MMU-5632684, R-MMU-5635838), and mitochondrial activity (R-MMU-5389840, R-MMU-611105, R-MMU-163210, R-MMU-8949215).

#### Generation of cis-co-accessibility networks (CCANs)

We used the Cicero^47^ package (v1.3.6) to generate *cis*-co-accessibility maps of regulatory elements and gene promoters. Young and aged MuSCs were separately reclustered using the peak matrix from the MuSC cluster and UMAP embeddings were generated for each group. The peak matrix for each group was binarized and converted to cell_data_set objects using the ‘make_atac_cds’ function in Cicero. Peaks containing at least one Tn5 insertion were identified and size factors were estimated using the ‘detect_genes’ and ‘estimate_size_factors’ functions in Monocle3^100^. The resulting objects were combined with the previously determined UMAP coordinates to create CDS objects for Cicero using the ‘make_cicero_cds’ function in Cicero. The number of cells per aggregate (‘k’ parameter) was selected as the maximum number of cells such that the median number of cells shared between aggregates was 0 and no more than 10% of cells were shared between paired aggregates on average (k=35 for young, k=20 for aged). This yielded 232,035 and 321,502 pairs of accessible sites with positive co-accessibility scores in young and aged MuSCs, respectively, which we filtered using a co-accessibility score of 0.1. Next, we generated the *cis*-co-accessibility maps using the ‘run_cicero’ function with window=5e5 and sample_num=100. Linked sites were annotated for genomic features using the ChIPseeker package (v1.30.3)^82^. We identified *cis*-co-accessibility networks (CCANs) in young and aged maps using the ‘generate_ccans’ function with a co-accessibility cutoff of 0.1, which maximized the number of CCANs across both maps. We used the maxmatching package (v0.1.0) in R to identify stable CCANs such that the total number of co-accessible elements between pairs of CCANs was maximized. To identify differences in connectivity within matched CCANs, we found CCANs in merged young and aged datasets (k=50) with the same parameters and found CCANs that matched CCANs matched between young and aged datasets.

#### Enrichment of CCANs within TADs

Pairs of *cis*-co-accessible sites were grouped by the distance between them into bins of 25kb. We calculated the fold enrichment of sites within TADs per distance bin as the ratio between the number of pairs within an individual TAD and the number of pairs that lay in different TADs. A null background comparison was performing this same analysis on sites that were shuffled in each distance bin and chromosome such that links were randomly assigned between sites. This procedure was repeated 100 times per distance bin.

#### Visualization of Peak-to-gene Linkages and Cicero Connections

Integrated visualizations of single cell ATAC tracks, peak-to-gene linkages, and *cis*-co-accessible sites were made using the Plotgardener^75^ package (v1.2.0). ATAC signal tracks were normalized to the number of reads in TSS regions to account for differences in sequencing depth and sample quality. The heights of linkages and *cis*-co-accessible sites are scaled to the distance between anchors.

## Figure and Table Captions

**Supplementary Figure 1. Quality control assessments of Hi-C datasets from young and aged muscle stem cells. A)** Pairwise Pearson correlation scatterplots of observed/expected counts in young and aged replicates at 250 kb resolution. **B)** Number of contacts at 250 kb resolution between loci of increasing genomic distance for each young and aged replicate compared with the merged replicates. **C)** Pie chart of the number of reads that map to inter- and intra-chromosomal interactions at 100 kb resolution. Long-cis and short-cis interactions refer to interactions on the same chromosome between loci separated by >20 kb and <20 kb, respectively. **D)** Overlapped first principal components of the Pearson correlation matrix of young and aged MuSC contact maps. **E)** Sizes of A/B compartments in young and aged MuSCs in log_10_(bases). **F)** Saddle plots of genome-wide A/B compartment enrichment. Eigenvector percentiles used to calculate compartment enrichment are shown below. **G)** Average RPKM signals per compartment switching region of ATAC-Seq and H3K4me3 datasets between aged and young MuSCs. **H)** Enriched GO and Reactome terms for genes in chromatin compartment groups identified through GSEA.

**Supplementary Figure 2. Definition and annotation of Topologically Associated Domains in young and aged muscle stem cells. A)** Optimization of the contact map resolution and FDR parameter used in HiCExplorer (hicFindTads) for finding TADs. **B)** Pearson correlation matrices of ATAC and H3K4me3 signals inside and within 15 kb of TAD domains. **C)** Distribution of CTCF motif enrichment across TAD domains. **D)** Comparison of the distribution of TAD boundaries across gene bodies. **E)** Annotation of TAD boundaries. **F)** Representative GO and Reactome terms for housekeeping genes enriched at shared TAD boundaries. **G)** Aggregate contact maps at lost, gained, and shared TAD boundaries in log_2_(observed/expected) counts. **H)** TAD separation scores and **I)** gene expression at lost, gained, and shared TAD boundaries. **J)** Distance in basepairs of TAD boundaries to the nearest expressed TSS. A vertical line is drawn at 1kb, representing the defined region for promoters. **K)** Diagram of intra-TAD connectivity. The mean read counts per intra-TAD interaction are divided by the sum of mean read counts for all interactions per TAD. Intra-TAD regions are highlighted in green while inter-TAD regions are highlighted in purple. **L)** Comparison of intra-TAD connectivity by compartment. **M)** ATAC and H3K4me3 signals in RPKM (mean +/- SEM) against binned intra-TAD connectivity percentiles. Statistical comparisons are pairwise Mann-Whitney comparisons between each percentile bin against the mean of all bins within each signal type and age group. (p-values: *<0.05, **<1e-2, ****<1e-3, ****<1e-4). **N)** Comparison of intra-TAD connectivity between unique and shared TADs. All statistical comparisons are unpaired Mann-Whitney U-tests.

**Supplementary Figure 3. Characterization of genomic interactions and chromatin loop domains in young and aged muscle stem cells. A)** Venn diagram of chromatin loops. **B)** Comparison of the distribution of loop anchors across gene bodies. **C)** Annotation of loop anchors. **D)** Distribution of CTCF motif enrichment between loop anchors. **E)** Classification of loop anchors across loop types. **F)** H3K4me3 and ATAC-seq signal strengths in RPKM (mean +/- SEM) at anchors of promoter-distal loops. Statistical comparisons are unpaired Mann-Whitney U-tests between age groups within each bin. **G)** Comparison of gene expression between or outside of chromatin loop anchors. Statistical comparisons are unpaired Mann-Whitney U-tests. **H)** Hierarchical clustering of inferred differentially bound transcription factors by overlap of TF binding sites. Motif names are colored by the direction of enrichment (Young—blue; Aged—red).

**Supplementary Figure 4. Quality assessments of single young and aged muscle stem cell multi-omic datasets. A)** QC plots of TSS enrichment against the log_10_(number of unique fragments) per cell across samples. **B) and C)** Density plots of **B)** fragment size and **C)** enrichment of insertion sites around TSS sites. Insets show data specific for MuSCs. **D)** Dot plots of gene activity scores (top) and integrated single cell RNA (scRNA) gene expression (bottom) for marker genes in identified cell type clusters. **E)** QC plots of previously collected MuSC scRNA datasets and age-matched datasets from Tabula Muris Senis (TMS). **F)** UMAP embeddings of scRNA data colored by cell type and split by dataset. **G)** Dot plot of marker gene expression in integrated scRNA datasets. **H)** Row-scaled heatmap of differentially accessible peaks in each cell type cluster.

**Supplementary Figure 5. Mapping gene expression regulatory networks in single young and aged muscle stem cells. A)** Annotation of union MuSC peakset and peaks in each peak-to-gene cluster. **B)** Histogram of the number of TADs per peak-to-gene linkage. **C)** Scatter plot of all identified co-accessible sites using Cicero. The minimum co-accessibility score threshold (0.1) used to filter sites is shown in red. **D)** Intra-TAD enrichment (log2 odds ratio) of linked peaks compared to randomly linked distance-matched peaks across TADs across different co-accessibility thresholds. **E)** Numbers of sites in matched and unmatched CCANs. **F)** Average fractions of CCAN members that leave, join, or remain in matched CCANs during aging. **G)** Expression of differentially expressed genes in matched CCANs. **H) and I)** Representative plots showing altered connectivity within young and aged CCANs relative to CCANs identified in merged young and aged data for the **H)** *Mta2* and **I)** *Ndufc1* loci. The top group of plots shows the CCAN containing each gene. The bottom group shows zoomed-in connections in each CCAN that lie in each gene body which is marked by a green rectangle.

**Supplementary Table 1. Quality assessment metrics of single cell ATAC datasets**.

