## Supplemental Figures and Tables for "Three-dimensional chromatin re-organization during muscle stem cell aging"

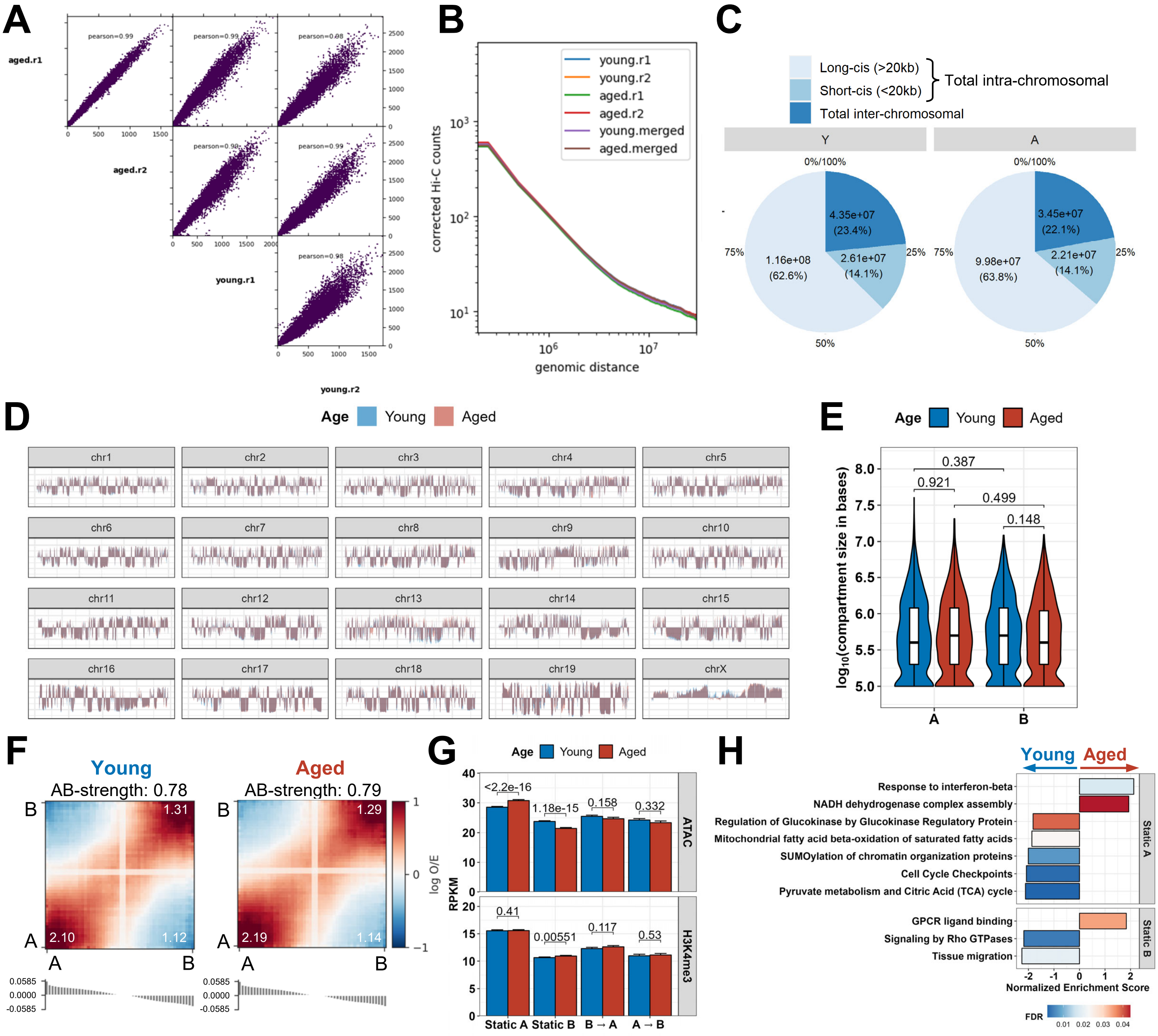


### Supplementary Figure 1

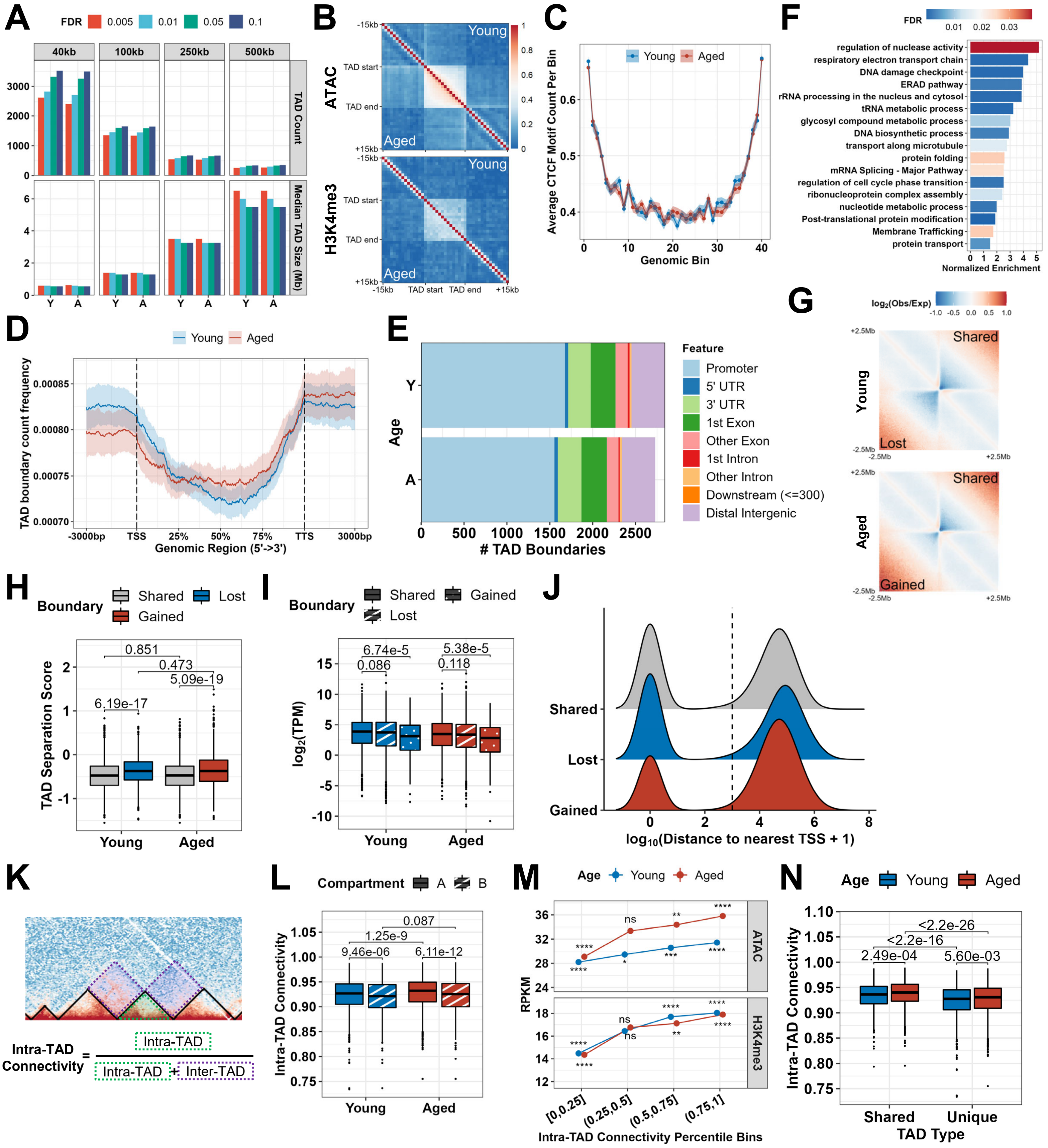


### Supplementary Figure 2

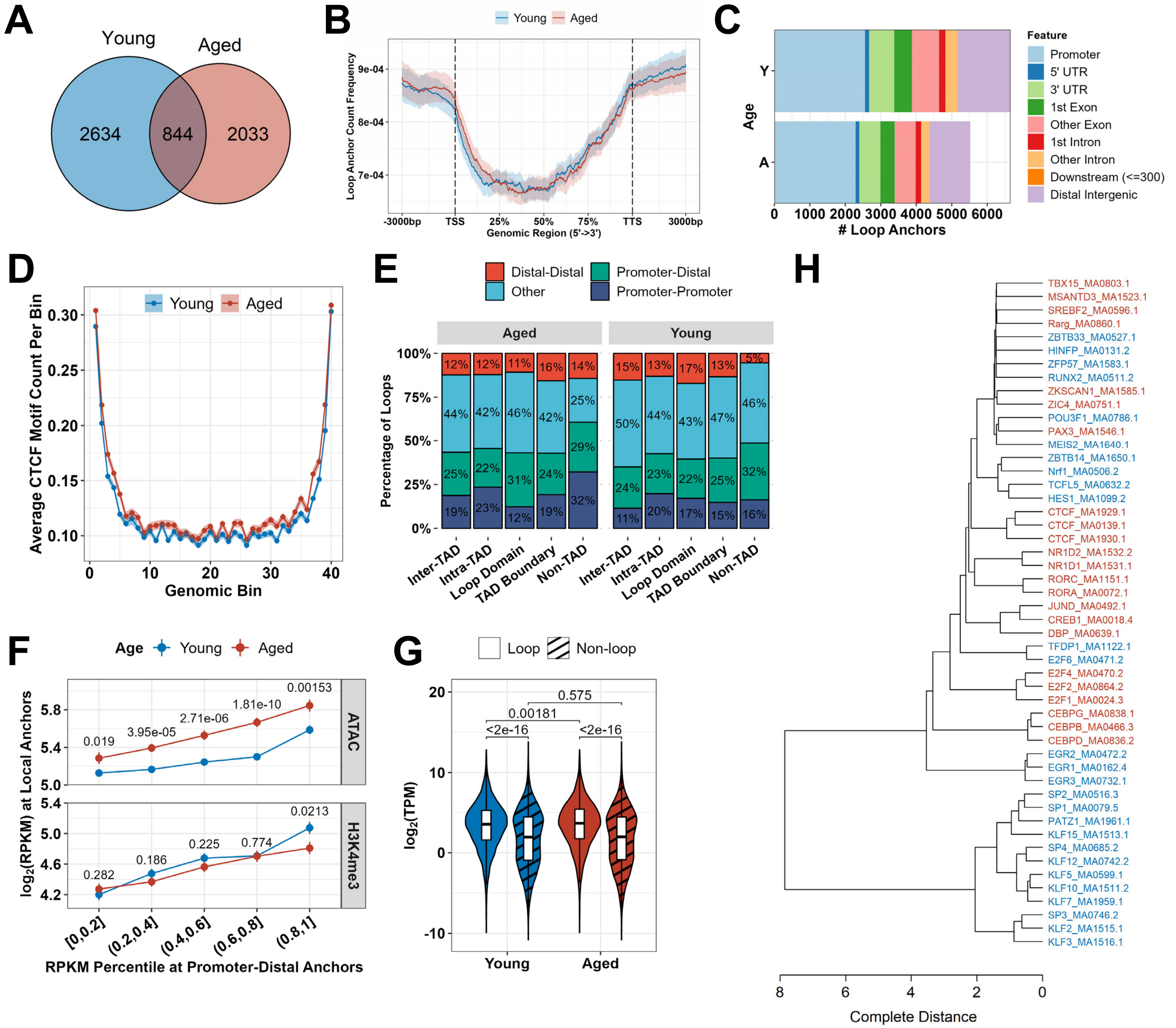


### Supplementary Figure 3

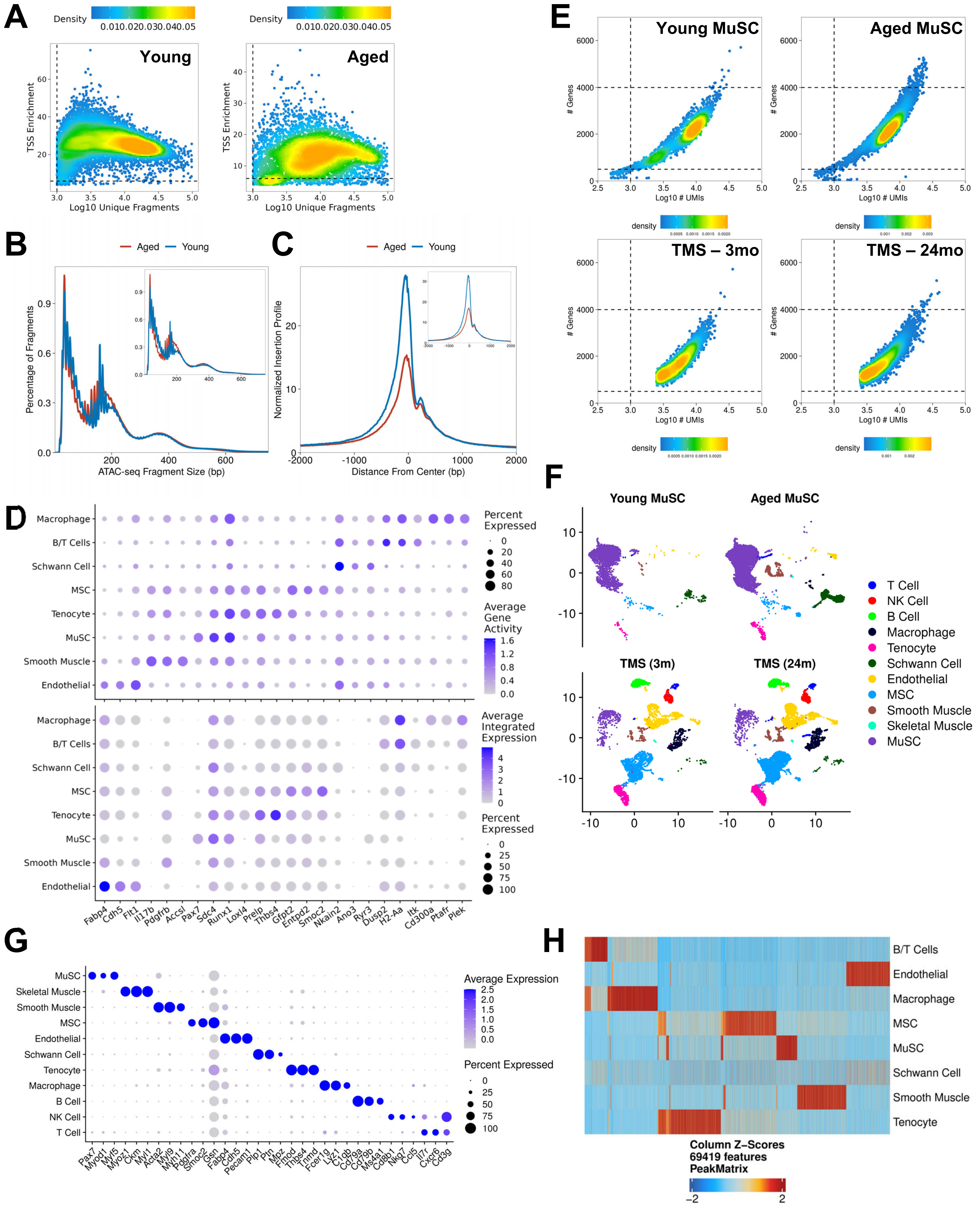


**Supplementary Figure 4**

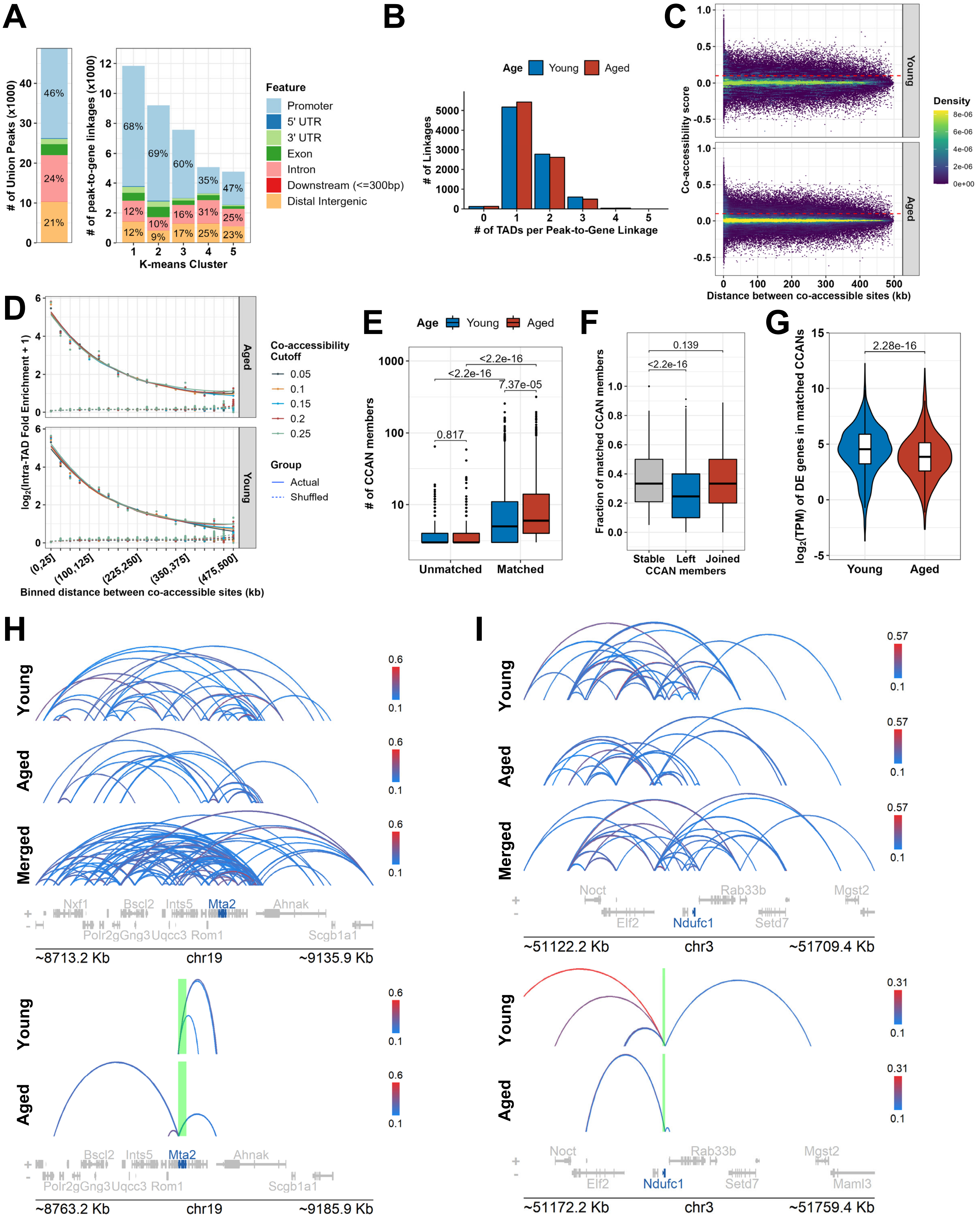


### Supplementary Figure 5

| Sample | # QC'ed Cells | Median Peaks per Cell | Median Fragments per Cell | Median TSS Enrichment | Median Blacklist Ratio | Median FRiP |
|---|---|---|---|---|---|---|
| Young | 10,705 | 2,344.5 | 5,426 | 25.1 | 0.0228 | 0.533 |
| Aged | 4,558 | 1,934.5 | 8,006 | 13.7 | 0.0210 | 0.271 |

| Sample | # QC'ed MuSCs | Median Peaks per Cell | Median Fragments per Cell | Median TSS Enrichment | Median Blacklist Ratio | Median FRiP |
|---|---|---|---|---|---|---|
| Young | 402 | 1,626 | 3,100 | 28.2 | 0.0248 | 0.518 |
| Aged | 178 | 6,316 | 23,004 | 15.8 | 0.0232 | 0.308 |

**Supplementary Table 1. Quality assessment metrics of single cell ATAC datasets.**
